# Shisa7-dependent regulation of GABA_A_ receptor single-channel gating kinetics

**DOI:** 10.1101/2022.02.11.480135

**Authors:** David Castellano, Kunwei Wu, Angelo Keramidas, Wei Lu

## Abstract

GABA_A_ receptors (GABA_A_Rs) mediate the majority of fast inhibitory transmission throughout the brain. Although it is widely known that pore-forming subunits critically determine receptor function, it is unclear whether their single-channel properties are modulated by GABA_A_R-associated transmembrane proteins. We previously identified Shisa7 as a GABA_A_R auxiliary subunit that modulates the trafficking, pharmacology, and kinetic properties of these receptors. In particular, Shisa7 accelerates receptor deactivation; however, the underlying mechanisms by which Shisa7 controls GABA_A_R kinetics have yet to be determined. Here we have performed single-channel recordings and find that while Shisa7 does not change channel slope conductance, it reduces the frequency of openings. Importantly, Shisa7 modulates GABA_A_R gating by decreasing the duration and open probability (P_o_) within bursts. Kinetic analysis of dwell times, activation modeling, and macroscopic simulations indicate that Shisa7 accelerates GABA_A_R deactivation by governing the time spent between close and open states during gating. Together, our data provide a mechanistic basis for how Shisa7 controls GABA_A_R gating kinetics and reveal for the first time that GABA_A_R single-channel properties can be modulated by an auxiliary subunit. These findings shed light on processes that shape the temporal dynamics of GABAergic transmission.

## Introduction

GABA_A_Rs are pentameric ligand-gated ion channels assembled from 19 different pore-forming subunits (Olsen and Sieghart, 2009) These receptors are critical for providing fast inhibitory transmission throughout the brain (Farrant and Nusser, 2005; Mody and Pearce, 2004). Each subunit exhibits a similar topology (Olsen and Sieghart, 2009; Sigel and Steinmann, 2012) comprising four, helical transmembrane domains (TMD, M1-4), extracellular domains (ECD; pre-M1, M2-M3 linker, post-M4), and intracellular domains (ICD; M1-M2 and M3-M4 linkers). Following their heteropentameric assembly, receptors are trafficked to the cell surface and are distributed to either synaptic or extrasynaptic regions where they contribute to phasic and tonic inhibition, respectively (Farrant and Nusser, 2005; Lorenz-Guertin and Jacob, 2018; Martenson et al., 2017).

GABA_A_Rs harbor distinct binding sites and serve as targets for several endogenous ligands and drug classes including benzodiazepines, barbiturates, neurosteroids, general anesthetics, and alcohol (Castellano et al., 2021; Kim and Hibbs, 2021; Olsen, 2018; Scott and Aricescu, 2019). Upon agonist binding, GABA_A_Rs undergo a series of conformational changes that promote channel opening and increase chloride conductance. This process involves inter-subunit interactions and the rotation of subunit ECDs, engagement of structural elements located at the ECD-TMD interface, and displacement of the M2 helix lining the channel pore (Kim and Hibbs, 2021; Scott and Aricescu, 2019). Insights into the mechanisms that regulate GABAAR activity are fundamental towards understanding their functional role at inhibitory synapses.

The time course of synaptic currents is largely dependent on the kinetic behavior of individual receptors (Dixon et al., 2014; Keramidas and Harrison, 2010; Lester et al., 1990). Single-channel, patch-clamp recordings in various neuronal culture preparations (Brickley et al., 1999; Lorez et al., 2000; Macdonald et al., 1989; MacDonald et al., 1989; Twyman et al., 1990) and in heterologous cells expressing receptor subtypes (Dixon et al., 2014; Keramidas and Harrison, 2010, 2008; Lema and Auerbach, 2006) has provided much information detailing the importance of pore-forming subunits towards regulating GABA_A_R conductance and gating. However, the emergence of recently identified GABA_A_R-associated, transmembrane accessory proteins brings into question whether they also modulate the biophysical characteristics of these receptors.

Native receptor complexes are comprised of pore-forming subunits together with auxiliary subunits that alter channel expression and function (Castellano et al., 2021; Han et al., 2020; Maher et al., 2017; Yan and Tomita, 2012). Several GABA_A_R-associated, transmembrane accessory proteins have been recently discovered, which include Lhfpl4 (also known as GARLH4), Clptm1, and Shisa7 (Davenport et al., 2017; Ge et al., 2018; Han et al., 2019; Wu et al., 2018; Wu et al., 2021; Yamasaki et al., 2017). Specifically, our previous studies have characterized Shisa7 as a GABA_A_R auxiliary subunit that controls the trafficking, pharmacological, and kinetic properties of GABA_A_Rs (Han et al., 2019; Wu et al., 2021). Importantly, we found that Shisa7 accelerates deactivation kinetics of GABA_A_Rs and global deletion of Shisa7 slows IPSC decay in hippocampal neurons (Han et al., 2019), indicating a critical role of Shisa7 in modulating the temporal dynamics of inhibitory transmission. However, the mechanisms underlying how Shisa7 speeds up GABA_A_R deactivation remain unknown.

In this study, we sought to investigate whether Shisa7 modulates GABA_A_R single-channel properties. We find that while Shisa7 does not change GABA_A_R slope conductance, it reduces the frequency of opening events. Single-channel kinetic analysis reveals that Shisa7 alters the duration, P_o_, and mean close and open times within GABA_A_R bursts, specifically, by changing the time, but not proportion, spent in specific close and open states. Furthermore, Shisa7 reduces gating efficacy by increasing occupancy of the final close state preceding channel openings. Lastly, simulation of GABA currents demonstrates that Shisa7 can accelerate GABA_A_R deactivation through its alterations in single-channel burst duration and P_o_. Together our findings show that Shisa7 is a GABA_A_R auxiliary subunit capable of regulating single-channel gating kinetics and highlight the importance of understanding how auxiliary subunits can control the gating process of native GABA_A_Rs.

## Results

### Shisa7 modestly reduces GABA potency

We previously demonstrated that Shisa7 did not cause a significant change in GABA potency for α2β3γ2L GABA_A_Rs using concentrations ranging from 0.3 μM to 1 mM GABA (Han et al., 2019). Since a decrease in GABA potency may partially account for how Shisa7 accelerates deactivation, we re-performed GABA concentration-response experiments with a broader range of GABA concentrations (0.1 μM to 10 mM GABA) and used a faster solution exchange perfusion step system with lifted HEK293T cells expressing α2β3γ2L with or without Shisa7. We find that the peak amplitude elicited by 10 mM GABA is significantly higher in GABA_A_Rs with the presence of Shisa7 compared to control (Figure 1A-B), consistent with our previous findings (Han et al., 2019; Wu et al., 2021). Under these new conditions, we find that Shisa7 causes a very modest, yet significant right-shift in the GABA-concentration response curve (Figure 1C), suggesting that Shisa7 elicits a slight reduction in GABA potency in α2β3γ2L GABA_A_Rs.

**Figure 1.**
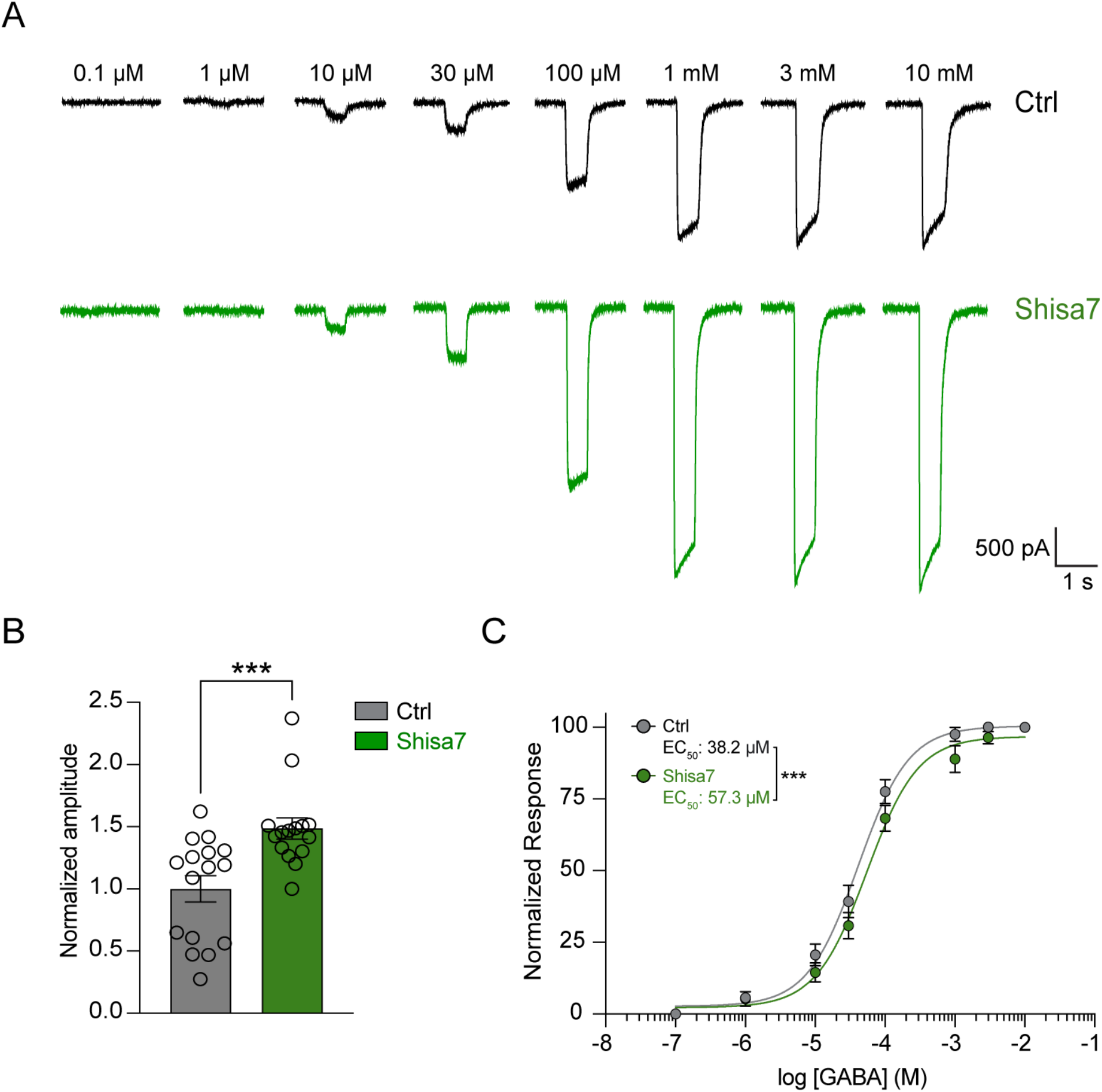
Shisa7 modestly reduces GABA potency. (**A**) Representative traces of GABA-evoked, whole-cell currents in HEK293T cells co-transfected α2β3γ2L GABA_A_Rs with pIRES2-EGFP (Ctrl; top) or pIRES2-Shisa7-EGFP (Shisa7; bottom). (**B**) Normalized amplitude responses demonstrating that Shisa7 increases GABA-evoked currents compared to control in response to 10 mM GABA (Ctrl, *n* = 16; Shisa7, *n* = 15; Mann-Whitney U, two-tailed). Error bars indicate S.E.M. (**C**) GABA concentration-response (0.1 μM -10 mM GABA) curves demonstrating that Shisa7 slightly reduces GABA potency in α2β3γ2L GABA_A_Rs compared to Ctrl. Data was normalized to the maximal response within the same cell (Ctrl, *n* = 16, LogEC_50_ = -4.42, EC_50_ = 38.2 μM ; Shisa7, *n* = 15, LogEC_50_ = -4.242, EC_50_ = 57.3 μM). Normalized data were fit by nonlinear regression. Data represents mean ± S.E.M. ***p < 0.005.

The shape of macroscopic GABA currents (I_GABA_) is determined by the number of receptors present at the surface (N), their P_o_, and GABA_A_R conductance (i_GABA_). Since Shisa7 increases peak GABA-evoked whole-cell currents, which is mostly due to increased surface trafficking (Han et al., 2019, Wu et al., 2021), yet only slightly reduces GABA potency, we set out to investigate whether Shisa7 alters GABA_A_R single-channel properties such as i_GABA_ or P_o_ and whether these effects could explain how Shisa7 accelerates deactivation (Han et al., 2019). We focused on one of the major synaptic GABA_A_R subtypes, α2β3γ2L, and performed single-channel, patch-clamp recordings under sub-saturating (3 µM) and saturating (10 mM) GABA conditions.

### Shisa7 does not change GABA_A_R slope conductance

First, we performed cell-attached, single-channel recordings of α2β3γ2L GABA_A_Rs in HEK293 cells in the presence or absence of Shisa7 to determine whether Shisa7 changes i_GABA_. A sub-saturating concentration of 3 µM GABA was included in the patch pipette, which elicits GABA_A_R single-channel opening and closing events (Figure 2A). After the patch baseline current stabilized, the pipette holding potential was changed in 25 mV increments from +100 to -100 mV (Figure 2A-B) and opening amplitudes were measured at each voltage step (Figure 2B). We find that α2β3γ2L GABA_A_Rs exhibits a slope conductance of 25 ± 0.5 pS (Figure 2C), similar to previous observations of other heteropentameric GABA_A_Rs subtypes (Dixon et al., 2014; Lema and Auerbach, 2006). In the presence of Shisa7, the slope conductance is 23 ± 0.3 pS, which is not significantly different compared to control (Figure 2C). Thus, although Shisa7 regulates GABA_A_R trafficking, pharmacology, and channel kinetics, it does not affect the slope conductance of α2β3γ2L receptors.

**Figure 2.**
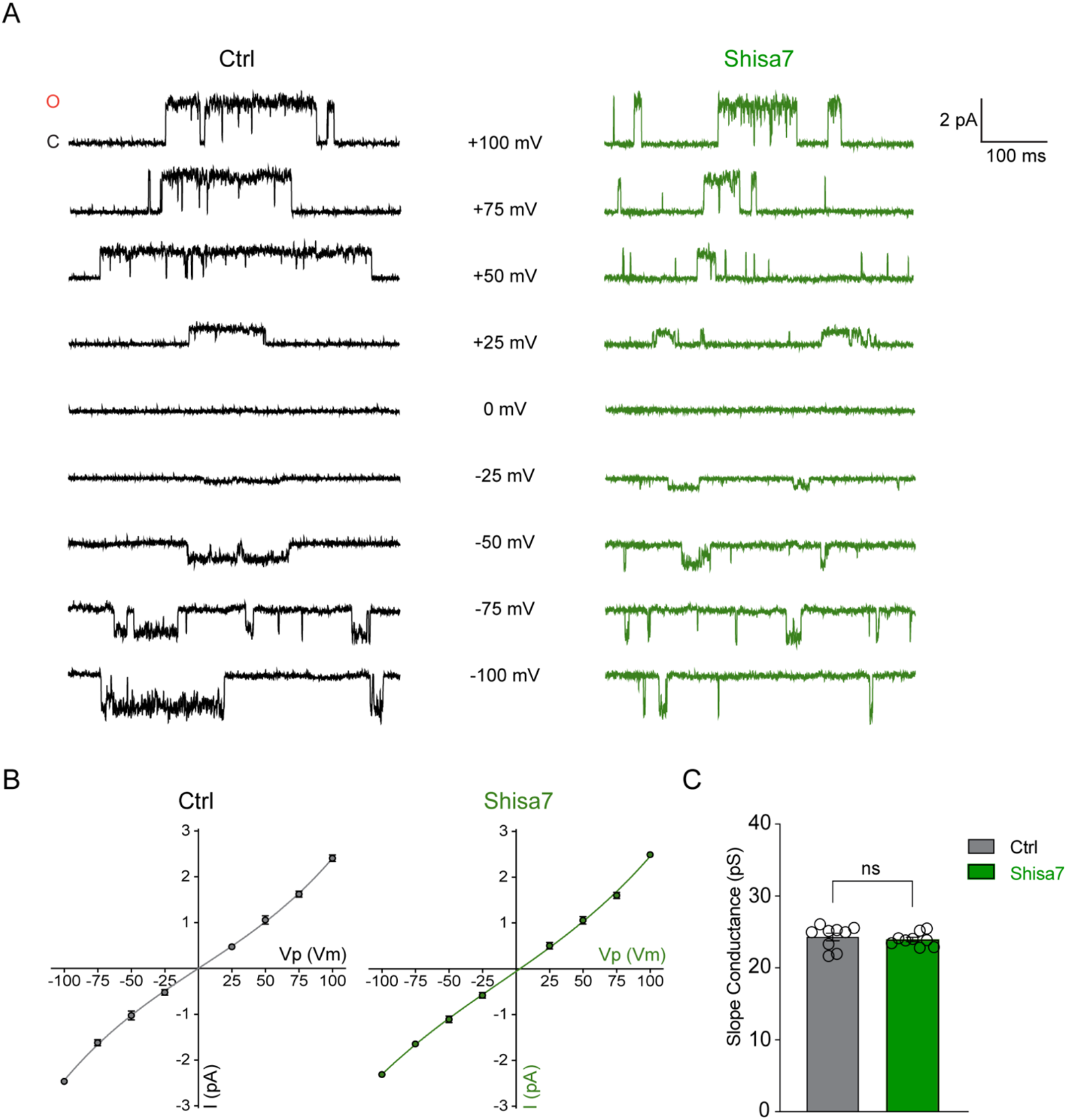
Shisa7 does not change GABA_A_R slope conductance. **(A)** Representative traces of cell-attached recordings of α2β3γ2L GABA_A_Rs co-transfected with either Ctrl or Shisa7. 3 μM GABA was included intra-pipette and the amplitude of single-channel openings were recorded at different membrane potentials ranging from +100 to -100 mV. **(B)** Plot depicting the current-voltage relationship between α2β3γ2L co-transfected with either Ctrl (*n* = 9) and Shisa7 (*n* = 9). **(C)** Bar graph depicting the α2β3γ2L GABA_A_R slope conductance between Ctrl and Shisa7 (Mann-Whitney U-test, two-tailed; p = 0.49). Vp = patch potential. Error bars indicate S.E.M.

### Shisa7 reduces the frequency of GABA_A_R single-channel opening events

Under sub-saturating GABA concentrations, GABA_A_R single-channel opening events present as either brief, isolated openings or bursts of openings (Dixon et al., 2014; Lema and Auerbach, 2006; Mortensen and Smart, 2007). To examine whether Shisa7 alters single-channel opening frequency, we quantified the number of openings from both event types using cell-attached recordings from cells exposed to 3 μM GABA at a holding potential of +100 mV. In α2β3γ2L GABA_A_Rs, both control and Shisa7 display single-channel currents with an amplitude of ∼2 pA at this holding potential (Figure 3A). We observe that Shisa7 strongly reduces the frequency of all single-channel opening events by ∼65% (Figure 3B). Further analysis of these events shows that Shisa7 reduces both isolated (Figure 3C) and burst opening (Figure 3D) events by 64% and 69%, respectively. Together, these results demonstrate that Shisa7 reduces the frequency of single-channel openings in α2β3γ2L GABA_A_Rs under sub-saturating conditions.

**Figure 3.**
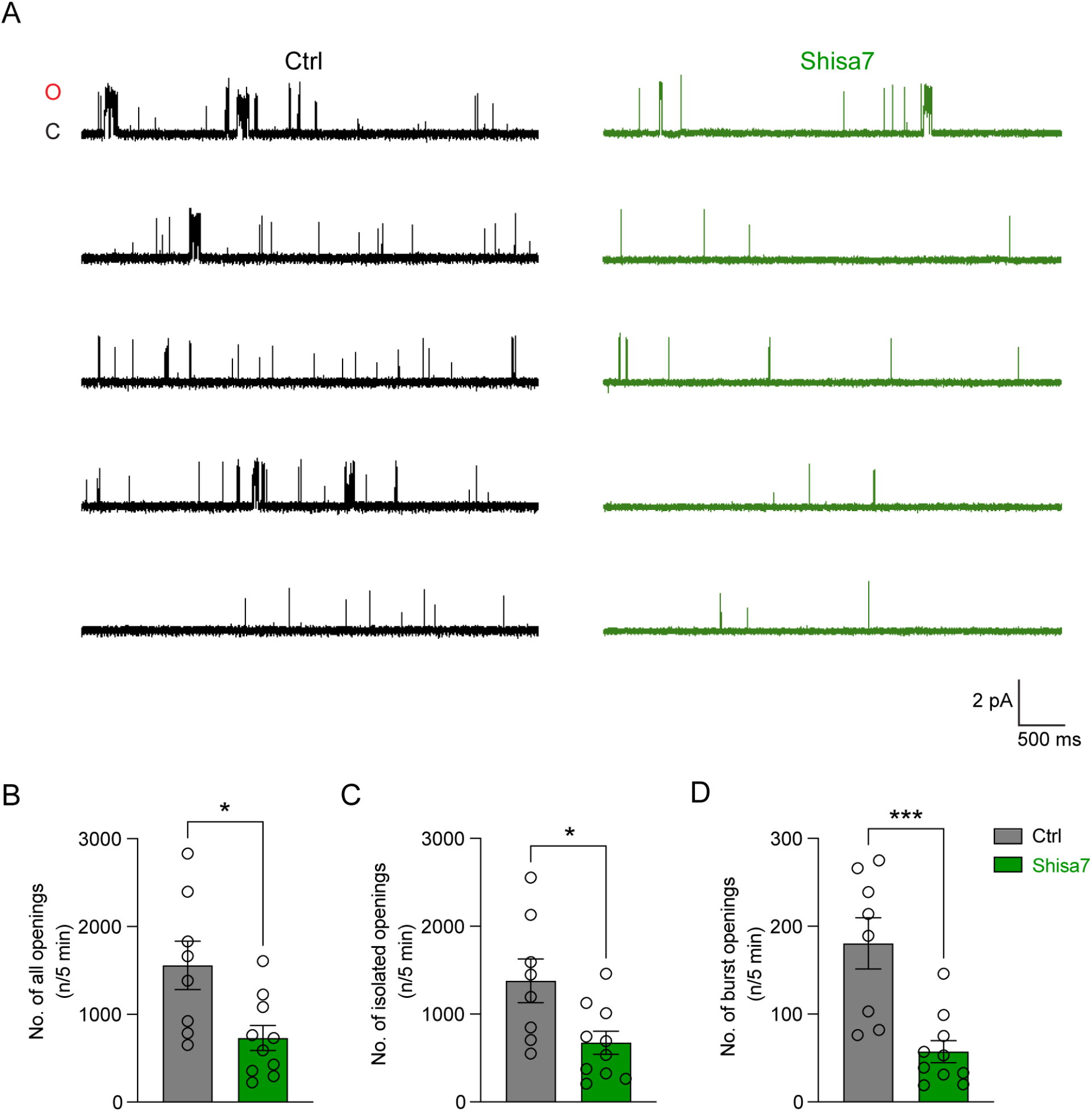
Shisa7 reduces the frequency of GABA_A_R single-channel opening events. **(A)** Representative traces of cell-attached recordings of α2β3γ2L GABA_A_Rs co-transfected with either Ctrl or Shisa7 in the presence of 3 μM GABA intra-pipette and held at +100 mV (O, open; C, close). **(B-D)** Bar graphs depicting the frequency of all GABA_A_R opening events (**B**), including isolated openings (**C**) and burst openings (**D**) (Ctrl, *n* = 8; Shisa7, *n* = 10, Mann-Whitney U-test, two-tailed). *p < 0.05; ***p < 0.005. Error bars indicate S.E.M.

### Shisa7 decreases GABA_A_R burst duration and P_o_

We have also previously shown that Shisa7 can accelerate GABA_A_R deactivation kinetics (Han et al., 2019). However, a mechanistic basis for this phenomenon has not been identified. At the microscopic level, it has been demonstrated that GABA_A_R deactivation kinetics are largely controlled by the duration and P_o_ of bursts (Dixon et al., 2014; Jones and Westbrook, 1995). To determine whether Shisa7 regulates single-channel burst kinetics, we analyzed burst activity elicited by 10 mM GABA in cell-attached configuration. Given that two GABA binding sites reside on GABA_A_Rs (Baumann et al., 2003; Masiulis et al., 2019), full occupancy of these sites using saturating GABA concentration enables us to negate the effects of agonist binding during the gating process (Lema and Auerbach, 2006; Mortensen and Smart, 2007). We performed maximum likelihood (MIL) fitting to obtain critical shut time (*τ*_crit_) values that were used to identify individual bursts. The *τ*_crit_ value was determined for each patch to preserve the 3 shortest close (C) and 3 shortest open (O) states that were consistent across patches. In α2β3γ2L GABA_A_Rs, we observe at least 3 modes of burst activity based on P_o_ (Figure 4A), as described in previous studies (Brodzki et al., 2020; Dixon et al., 2014; Lema and Auerbach, 2006) in both control and Shisa7 conditions. As we were interested in understanding how burst kinetics determine GABA_A_R deactivation, our analyses pooled all observed modes of bursts together. We find that Shisa7 decreases the activation of α2β3γ2L GABA_A_Rs by reducing both the burst duration and P_o_ (Figure 4B). Furthermore, Shisa7 increases the burst mean close time while also reducing burst mean open time (Figure 4B). Together, these findings demonstrate that in addition to decreasing opening frequency, Shisa7 can alter GABA_A_R single-channel burst properties.

**Figure 4.**
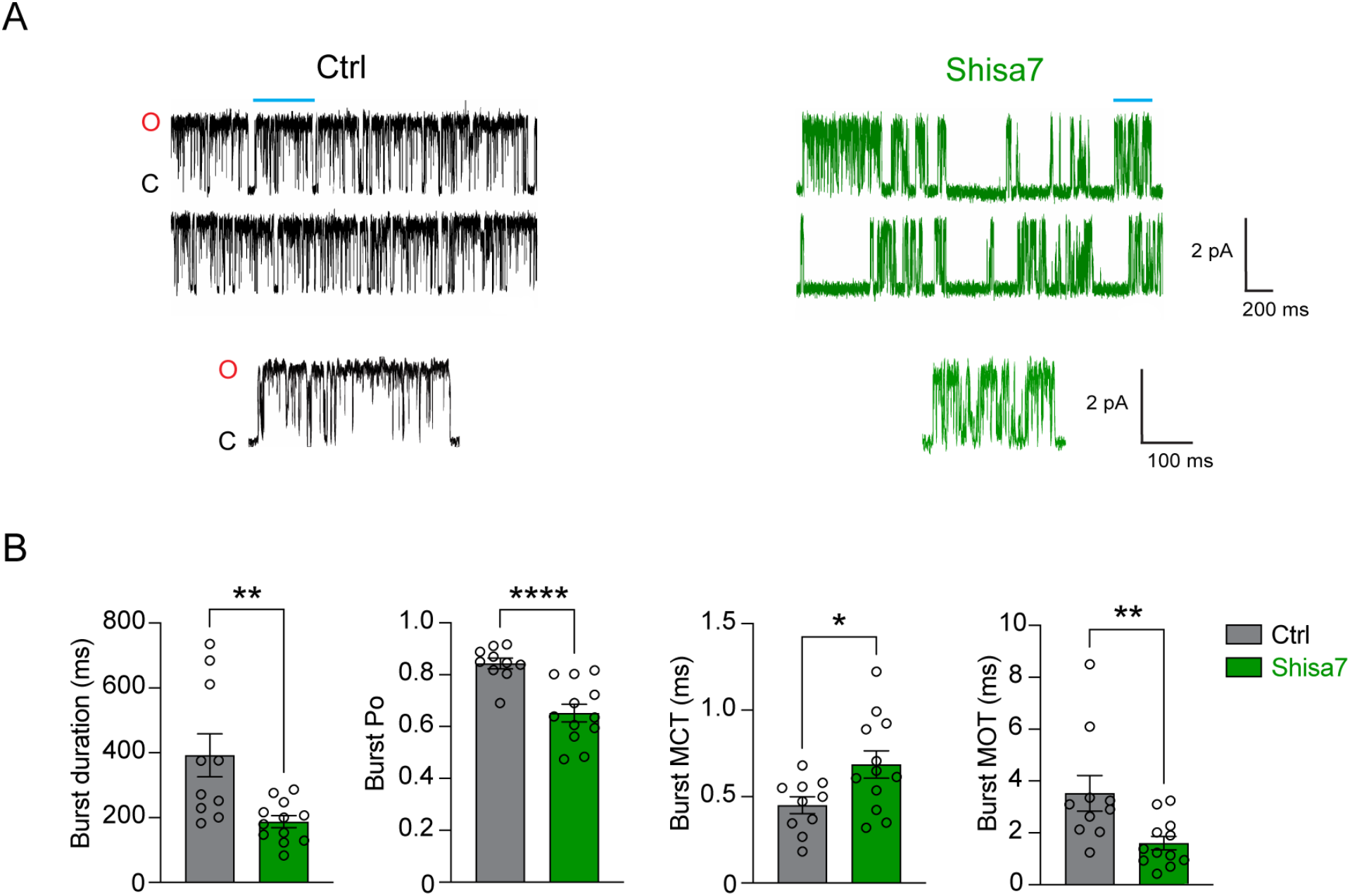
Shisa7 decreases GABA_A_R burst duration and P_o_. **(A)** Representative traces of single-channel clusters of bursts elicited by 10 mM GABA in α2β3γ2L GABA_A_R co-transfected with either Ctrl or Shisa7 (top), with the blue bar indicating representative bursts from both conditions (bottom). **(B)** Bar graphs depicting mean duration, mean P_o_, mean close time (MCT), and mean open time (MOT) of single-channel bursts (Ctrl, *n* = 10; Shisa7, *n* = 12; Mann-Whitney U-test, two-tailed, *p = 0.05, **p = 0.005, ****p < 0.0001). Error bars indicate S.E.M.

### Shisa7 alters close and open dwell time components during bursts

Several linear models of GABA_A_R activation during bursts have been previously explored (Dixon et al., 2014, 2015; Keramidas and Harrison, 2010; Lema and Auerbach, 2006), utilizing different schematic arrangements of the 3C and 3O states. To further understand how Shisa7 modulates GABA_A_R single-channel burst kinetics, we selected a 3C3O model scheme (Figure 6A) that displayed the largest best fit (log likelihood (ΣLL)) value for each patch between control and Shisa7 conditions. This scheme has been used previously to model GABA_A_R activation during bursts (Dixon et al., 2014; Keramidas and Harrison, 2010; Lema and Auerbach, 2006).

Analysis of the three close components reveals that Shisa7 increases the time constants of the two shortest close states without affecting the proportion spent in any of the 3 close states (Figure 5A, 5C, 5D). In contrast, Shisa7 decreases the time constants of the two longest open states without also altering open state proportions (Figure 5B, 5E, 5F). Given that the time constants of the two shortest close states and two longest open states operate within micro- and millisecond timescales, respectively, the decrease in burst P_o_ by Shisa7 is most likely driven by the reduction in the duration of long open states, with a smaller contribution also arising from brief close states. Together, these results suggest that Shisa7 changes GABA_A_R burst properties by altering the dwell time constants operating in particular close and open states during the gating process.

**Figure 5.**
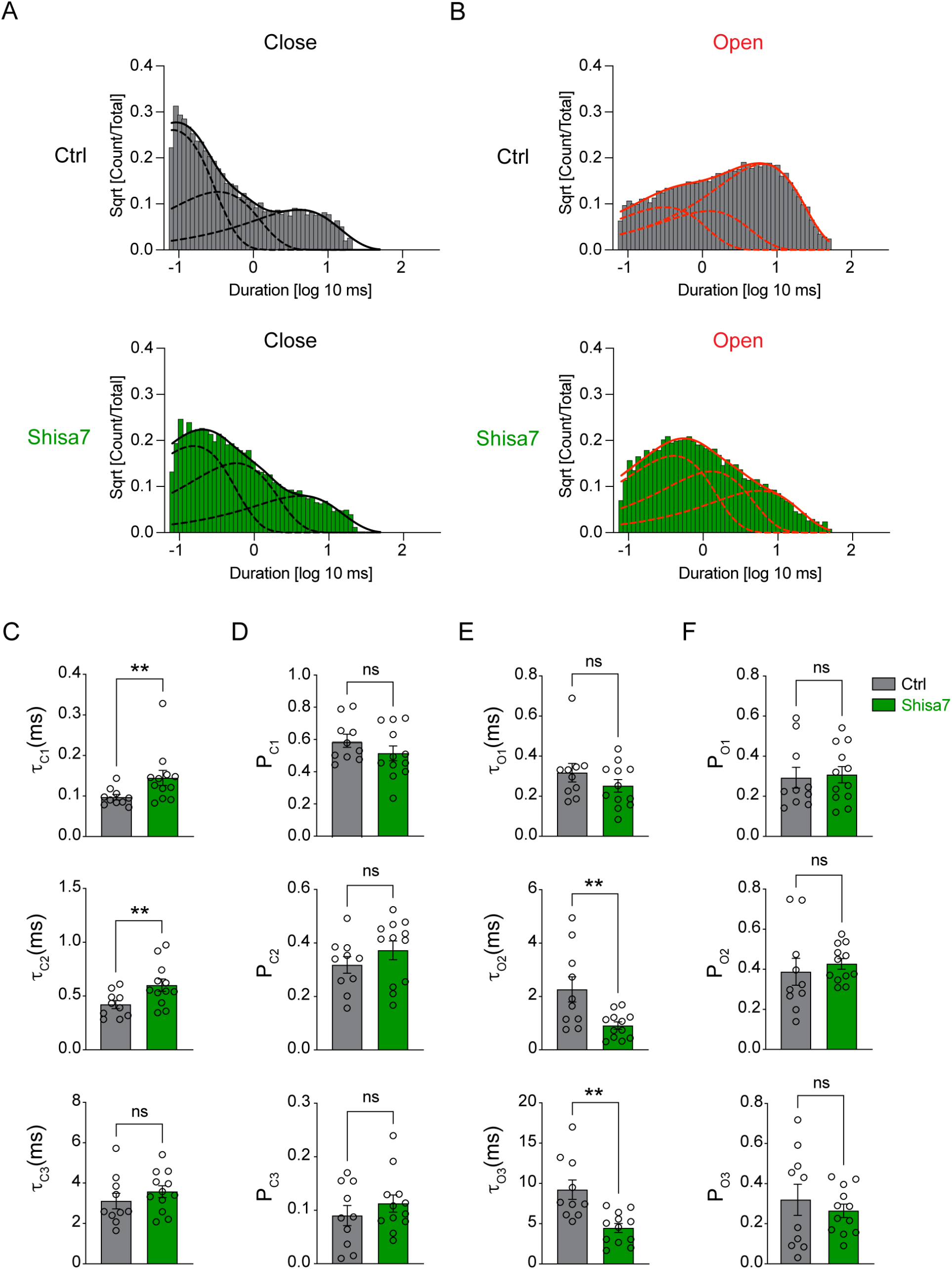
Shisa7 alters close and open dwell time components during bursts. **(A)** Close dwell time histograms of Ctrl (top) and Shisa7 (bottom) fitted with three exponential functions (black dotted lines) and the composite density function (black solid line). **(B)** Open dwell time histograms of Ctrl (top) and Shisa7 (bottom) fitted with three exponential functions (red dotted lines) and the composite density function (red solid line). **(C)** Mean time constants of the three close components. **(D)** Percentage of the three close components. **(E)** Mean constants of the three open components. **(F)** Percentage of the three close components. Ctrl, *n* = 10; Shisa7, *n* = 12; Mann-Whitney U-test, two-tailed, **p < 0.005. Error bars indicate S.E.M.

### Kinetic modeling of GABA_A_R bursts indicates that Shisa7 reduces the efficacy of channel opening

Next, we determined the forward and reverse rate constants between state transitions for each patch using the aforementioned 3C3O scheme arrangement (Figure 6A, Table 1). These were used to obtain equilibrium constants of state transitions within bursts. We find that compared to control, Shisa7 reduces E1 and E3, which represent transitions from the final close state directly proceeding channel opening states (Figure 6B, Table 2). We do not find significant differences between control and Shisa7 in the pre-activation step, ϕ, a short-lived, close-close state transition that ultimately progresses towards channel activation (Gielen et al., 2012, Kaczor et., 2021, Terejko et al., 2019, Dixon et al., 2014, Gielen et al., 2018, Lape et al., 2008) nor in Σ, a close-close transition that leads the channel further away from opening (Lema and Auerbach, 2008). Furthermore, there are no significant differences in E2 (Figure 6B), representing the occupancy of the shortest open state and is consistent with no changes in time nor proportion spent in the presence of Shisa7 for this state (Figure 5E, 5F). Together, these findings demonstrate that Shisa7 shortens α2β3γ2L GABA_A_R burst duration by reducing the efficacy of direct transitions from close-to-open states.

**Figure 6.**
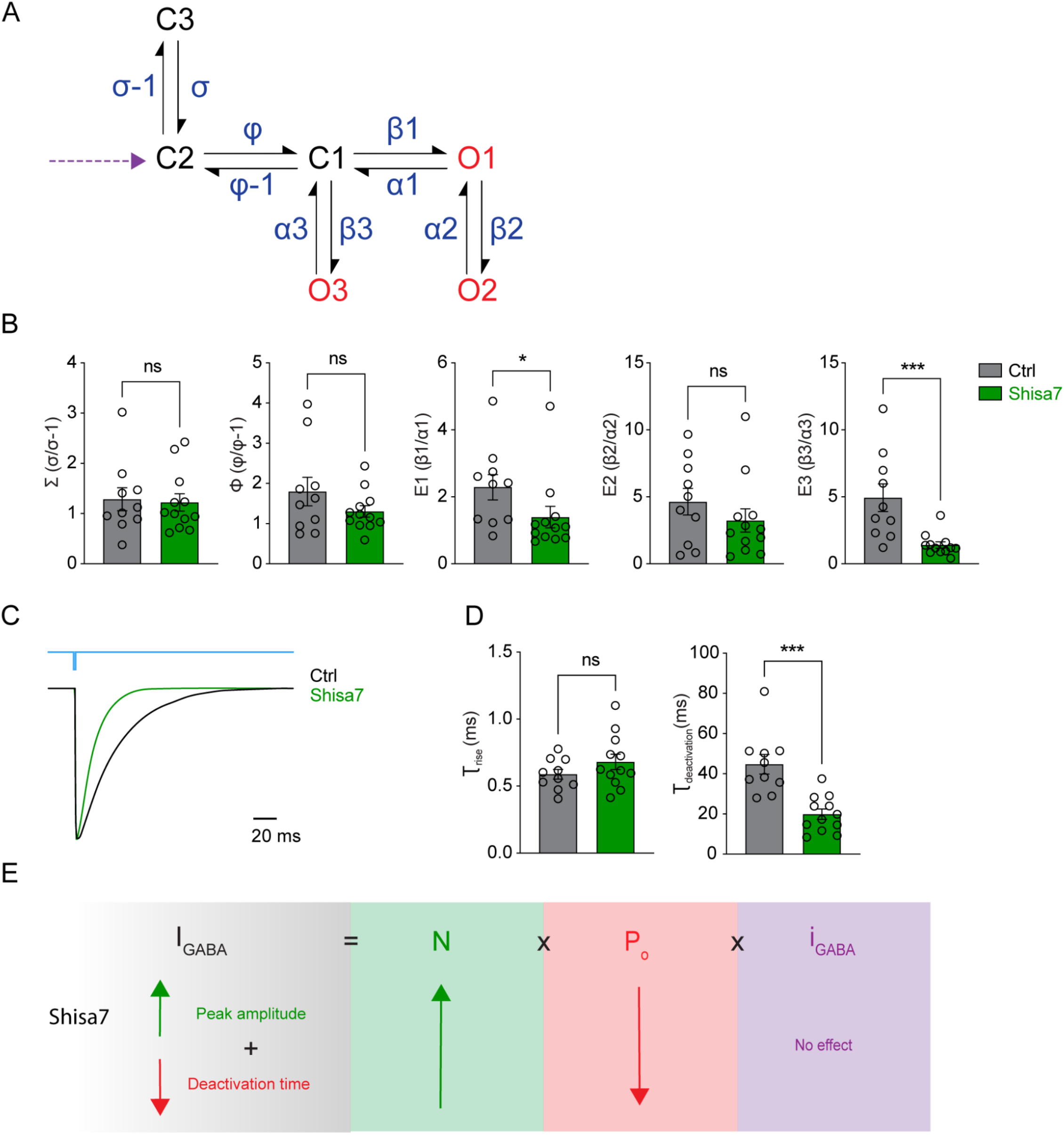
Kinetic modeling and simulation of GABA_A_R bursts reveal that Shisa7 increases occupancy of the final close state to accelerate receptor deactivation. **(A)** A linear, single-gateway model consisting of 3 close states (C1-3; black) and 3 open states (O1-3; red) was selected to model and simulate α2β3γ2L GABA_A_R burst activation. The rate constants (blue) depict the forward and reverse rate constants for each state transition. The purple arrow indicates GABA binding **(B)** Bar graphs depicting mean equilibrium constants between Ctrl and Shisa7. **(C)** A 1 ms stimulation pulse of 10 mM GABA (top) was used to simulate macroscopic GABA_A_R currents (bottom) using mean rate constants from Ctrl or Shisa7. **(D)** Bar graphs depicting representative simulated rise time (left) and weighted deactivation (right) between Ctrl and Shisa7. **(E)** A working model depicting Shisa7-dependent regulation of macroscopic GABA currents. Ctrl, *n* = 10; Shisa7, *n* = 12; Mann-Whitney U-test, two-tailed, *p < 0.05, ***p < 0.0005. Error bars indicate S.E.M. I_GABA_ = Macroscopic GABA current; N = Number of GABA_A_Rs present at surface; P_o_ = GABA_A_R open probability; i_GABA_ = GABA_A_R slope conductance.

**Table 1.** Rate constants for α2β3γ2L GABA_A_R bursts. Mean rate constants ± S.E.M for α2β3γ2L GABA_A_Rs co-transfected with GFP (Ctrl; *n* = 10) or Shisa7 (*n* = 12). Red indicates significant differences from Ctrl, *p < 0.05. Units are s^-1^. Mann-Whitney U-test, two-tailed, *p < 0.05.

|  | Rate Constants |  |  |  |  |  |  |  |  |  |
| --- | --- | --- | --- | --- | --- | --- | --- | --- | --- | --- |
| | $\sigma$ | $\sigma-1$ | $\varphi$ | $\varphi-1$ | $\beta 1$ | $\alpha 1$ | $\beta 2$ | $\alpha 2$ | $\beta 3$ | $\alpha 3$ |
| Ctrl | 474<br>$\pm 49$ | 482<br>$\pm 99$ | 2540<br>$\pm 279$ | 1787<br>$\pm 257$ | 7671<br>$\pm 1219$ | 4129<br>$\pm 1013$ | 1041<br>$\pm 293$ | 218<br>$\pm 45$ | 2912<br>$\pm 391$ | 818<br>$\pm 161$ |
| Shisa7 | 406<br>$\pm 36$ | 387<br>$\pm 60$ | 2066<br>$\pm 242$ | 1781<br>$\pm 240$ | 4982*<br>$\pm 918$ | 4748<br>$\pm 1171$ | 1229<br>$\pm 297$ | 403*<br>$\pm 62$ | 2272<br>$\pm 327$ | 2063*<br>$\pm 380$ |

**Table 2.**
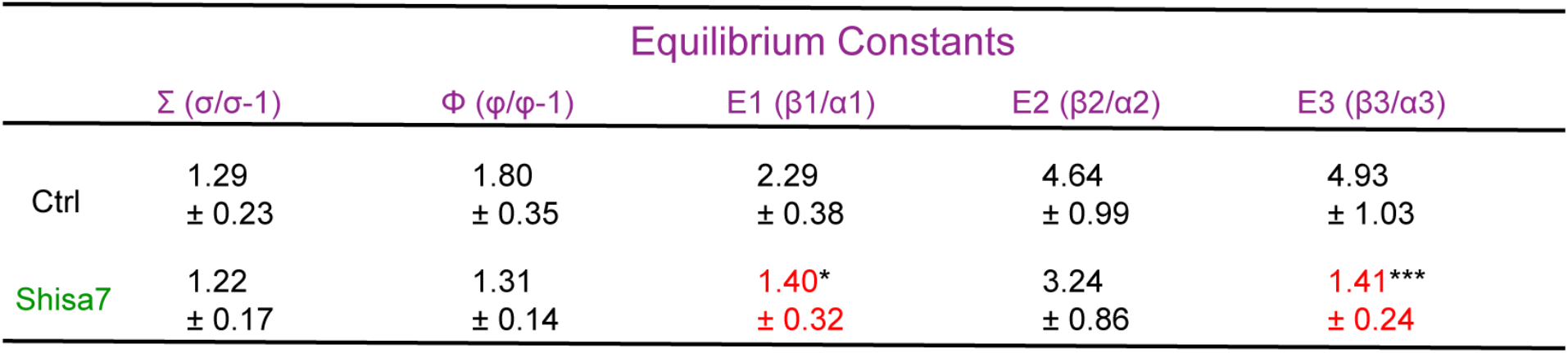
Equilibrium constants for α2β3γ2L GABA_A_R bursts. Mean equilibrium constants ± S.E.M for α2β3γ2L GABA_A_Rs co-transfected with GFP (Ctrl; *n* = 10) or Shisa7 (*n* = 12). Red indicates significant differences from Ctrl, *p < 0.05, ***p < 0.0005. Units are s^-1^. Mann-Whitney U-test, two-tailed, *p < 0.05, ***p < 0.0005.

### Model-based simulations reveal that Shisa7 accelerates synaptic current deactivation through changes in burst kinetics

Finally, we simulated synaptic currents using our kinetic model with averaged rate constants obtained from single-channel burst analyses. Our simulation protocol consisted of a 1 ms application of 10 mM GABA and used rate constants from each patch for both control and Shisa7 conditions (Figure 6C). We determined the time constants for the rise and deactivation time of simulated macroscopic current by fitting mono- and bi-exponential functions, respectively. We find that although Shisa7 does not change the rise time of the macroscopic current, it decreases the weighted deactivation time of the simulated GABA current (Figure 6D). Altogether, these findings demonstrate that changes to receptor burst gating kinetics mediated by Shisa7 are sufficient to recapitulate accelerated GABA_A_R deactivation as previously shown (Han et al., 2019).

## Discussion

Channel gating of ionotropic receptors plays a critical role in determining the strength and duration of synaptic transmission (Dixon et al., 2014; Keramidas and Harrison, 2010; Lester et al., 1990). In AMPA-type glutamate receptors, regulation of channel properties by transmembrane auxiliary subunits has been extensively investigated. Indeed, several auxiliary subunits such as the TARP and cornichon family have been shown to modulate AMPAR single-channel properties (Coombs and Cull-Candy, 2021; Jackson and Nicoll, 2011). In contrast, previous work aimed at characterizing GABA_A_R single-channel gating processes has mainly focused on differences between distinct receptor subtypes (Barberis et al., 2007; Dixon et al., 2014; Keramidas and Harrison, 2010, 2008; Lema and Auerbach, 2006; Macdonald et al., 1989; Twyman et al., 1990), critical domains involved in gating (Brodzki et al., 2020; Hernandez et al., 2017; Janve et al., 2016; Kash et al., 2003; Kisiel et al., 2018; Nors et al., 2021; Szczot et al., 2014; Terejko et al., 2019), and disease-related mutations in receptor subunits (Hernandez and Macdonald, 2019; Janve et al., 2016). Thus, it remained unknown whether GABA_A_R auxiliary subunits could directly modulate GABA_A_R single-channel properties.

We have recently shown that Shisa7, a single-pass transmembrane protein, interacts directly with GABA_A_R subunits and regulates both phasic and tonic GABAergic inhibition (Han et al., 2019; Wu et al., 2021). In addition to controlling GABA_A_R surface abundance, Shisa7 speeds up the deactivation kinetics of synaptic GABA_A_R subtypes such as α1β2γ2 and α2β3γ2 and loss of Shisa7 slows the decay of IPSCs in hippocampal neurons (Han et al., 2019). Interestingly, Shisa7 also enhances benzodiazepine potentiation of submaximal GABA responses, and global deletion of Shisa7 reduces the anxiolytic and sedative properties of diazepam in vivo (Han et al., 2019). While these findings demonstrate the importance of Shisa7 towards GABA_A_R function, how Shisa7 controls GABA_A_R kinetics are unclear.

We have now shown that while Shisa7 does not change single-channel conductance in α2β3γ2L GABA_A_Rs, it decreases the frequency of single-channel openings. Interestingly, single-channel recordings under saturating GABA conditions reveal that Shisa7 modulates GABA_A_R burst kinetic properties. Specifically, Shisa7 reduces burst duration, P_o_, and mean open time while also increasing mean close time. Further kinetic analyses of close and open dwell times reveal that Shisa7 changes the time, but not proportion, of certain states during gating. Determining the equilibrium constants between state transitions during gating suggests that Shisa7 decreases the ability to keep GABA_A_Rs open by increasing the occupancy of the final close state preceding openings, without changes to any direct close-close or open-open state transitions. Finally, simulating macroscopic currents from single-channel bursts demonstrates that Shisa7 accelerates GABA_A_R deactivation, consistent with our recent report (Han et al., 2019).

By using a faster solution-switching speed and a broader range of GABA concentrations, we also find that Shisa7 slightly decreases GABA potency. Although ligand potency is a function of both agonist binding and receptor gating (Colquhoun, 1998), we did not measure GABA binding affinity in the current study. It is worth noting that we observed a decrease in the frequency of channel opening events under sub-saturating GABA conditions. It is possible that this could be due to a combination of both decreased GABA binding affinity and receptor gating. Thus, future work is needed to determine whether Shisa7 can change binding affinities for GABA and other ligands at GABA_A_Rs. Previously, we have shown that Shisa7 enhanced whole-cell currents in heterologous cells expressing GABA_A_Rs (Han et al., 2019; Wu et al., 2021). Since Shisa7 does not affect slope conductance and only modestly changes GABA potency, our data indicate that the increased magnitude of peak, whole-cell GABA currents is most likely due to enhanced surface trafficking of GABA_A_Rs (Han et al., 2019; Wu et al., 2021).

Based on our data, we propose a working model describing Shisa7-dependent regulation of GABA_A_Rs in response to GABA activation (Figure 6E). At the macroscopic level, Shisa7 increases I_GABA_ due to enhanced N, as previously shown for both synaptic and extrasynaptic GABA_A_Rs via exocytosis (Han et al., 2019; Wu et al., 2021). At the single-channel level, we find that the presence of Shisa7 does not alter i_GABA_, indicating similar ionic conductance upon GABA_A_R activation. This suggests that the observed larger peak amplitude elicited by saturating GABA concentrations are primarily due to increased trafficking of GABA_A_Rs (Han et al., 2019; Wu et al., 2021) and not due to enhancement of Cl^-^ flux per receptor. Importantly, Shisa7 decreases burst P_o_ and duration, leading to faster GABA_A_R deactivation. Indeed, Shisa7 reduces channel openings primarily by shortening the duration of the two longest opening states during bursts and by increasing occupancy of the final close state. Taken together, in the presence of Shisa7 the activation of more receptors at the surface that gate more rapidly elicits both larger and faster GABA_A_R-mediated currents. These findings provide a mechanistic basis for Shisa7-dependent regulation of GABA_A_R gating at the single-channel level and shed new light on processes that control GABA_A_R activation.

## Materials and Methods

### Plasmids

Human α2 (NM_000807.3; pcDNA3.1-Zeo^+^), γ2L (NM_198903.2; pcDNA3.1-Zeo^+^), and Shisa7 (NM_001145176.2; pIRES2-EGFP) plasmids were generated by Genscript (GenScript, Piscataway, NJ). pIRES2-EGFP vector was a gift from Dr. Roger Nicoll’s lab at UCSF. Human β3 (NM_000814.6; pcDNA3.1-Zeo^+^) was a gift from Dr. Joseph Lynch’s lab at The University of Queensland, Australia.

### Cell Culture and Transfection

HEK293T cells (ATCC; CRL-11268) were maintained with culture media containing 1% penicillin-streptomycin (GIBCO), 10% FBS (GIBCO) in Dulbecco’s Modified Eagle’s Medium (DMEM, GIBCO), in a humidified incubator at 37 °C with 5% CO2. HEK293T cells were seeded onto eight 12 mm coverslips that were plated on 6 cm dish and transfected with human α2, β3, and γ2L GABA_A_R subunits with either pIRES2-EGFP or pIRES2-Shisa7-EGFP at a ratio of 1:1:3:3 (2 μg total cDNA), respectively. Plasmids were mixed in 94 μL 37°C-warmed DMEM without FBS and penicillin-streptomycin, then 6 μL FuGENE HD Transfection Reagent (Promega, USA) was added and vortexed to obtain a 3:1 reagent-to-DNA ratio. After 15 minutes to allow transfection reagent-DNA complex formation, the mixture was added dropwise to the cells.

### Whole-Cell Recordings

Whole-cell recordings were made from GFP-positive HEK293T transfected with α2β3γ2L GABA_A_Rs and either pIRES2-GFP or pIRES2-Shisa7-EGFP. The external solution contained (in mM): 140 NaCl, 5 KCl, 10 HEPES, 10 D-glucose, 1 MgCl_2_, 2 CaCl_2_ (pH 7.4, NaOH, ∼305-310 mOsm). The internal solution contained (in mM): 145 CsCl, 2 CaCl_2_, 2 MgCl_2_, 2 Mg-ATP, 10 HEPES and 10 EGTA, pH 7.4, CsOH, 305 mOsm). For GABA concentration-response curves, 1M GABA (Tocris) was prepared fresh from powder daily with the external solution and serial dilutions were made to obtain 8 GABA concentrations ranging from 0.1 μM to 10 mM. Single, isolated GFP-positive HEK293T cells were selected for concentration-response, whole-cell recordings. Borosilicate, filamented glass pipettes (OD: 1.5 mm, ID: 1.1 mm; Sutter Instrument) with an open tip resistance ranging from 3-6 MΩ were used. After obtaining whole-cell configuration, HEK293T cells were held at -40 mV and lifted to be positioned in front of a 3-barreled, square glass tube. During electrophysiological recordings, a VCS-8, computer-controlled valve system (Warner Instruments, USA) and VCS-77CSP8 fast perfusion stepper (Warner Instruments, USA) were used to control solution switching between two barrels: a control barrel containing external solution and a drug barrel containing GABA. A mini 8-to-1 manifold was used to load different GABA concentrations in the drug barrel. Whole-cell recording responses were collected with a Multiclamp 700B amplifier (Axon Instruments, USA) and digitized using Digidata 1400B (Molecular Devices, CA). Currents were sampled at 10 kHz and low-pass filtered at 2 kHz. Series resistance was monitored and not compensated, and cells that exhibited Rs variability by 25% during recording sessions were discarded.

### Single-Channel Recordings

Cell-attached, single-channel recordings were made from GFP-positive cells 24 hours post-transfection at room temperature (20–22°C). Current signals were acquired using an Axopatch 200B amplifier (Molecular Devices, USA) at a sampling rate of 100 kHz and 10 kHz Bessel low-pass filter. The data was digitized using a Digidata 1440B digitizer using pClamp 10 (Molecular Devices, USA). Coverslips containing transfected HEK293T cells were perfused with an external solution containing the following (in mM): 102.7 NaCl, 20 Na-gluconate, 2 KCl,14 D-glucose, 15 Sucrose, 10 HEPES, 20 TEA-Cl, 1.2 MgCl_2_, 2 CaCl_2_ (pH 7.4; NaOH; ∼320-330 mOsm). The internal solution was similar to the external solution, with the addition of either 3 μM or 10 mM GABA (Tocris; Cat. No. 0344). Sylgard 184 (Dow Corning, USA) was applied to the tip of each pipette, heated using a VT-750C Varitemp heat gun, and fire-polished using an MF-830 microforge (Narishige, Japan). After obtaining cell-attached mode, a pipette potential of +100 mV was applied. Borosilicate, filamented glass pipettes (OD: 1.5 mm, ID: 0.87 mm; Sutter Instrument) were pulled using a P-97 puller (Sutter Instruments, USA). The pipette tip resistance ranged from 10–25 MΩs.

### Single-Channel Kinetic Analysis

The QuB software was kindly provided by Dr. Gabriela Popescu’s lab at University at Buffalo, SUNY, and was used for kinetic analysis of single-channel bursts. Care was taken to analyze off patches that only displayed minimal overlapping opening events (<1% of total duration analyzed) at the beginning of the recording. Patches that exhibited ∼5000-15000 events were used for kinetic analysis. After correction for any baseline drift, single-channel data were idealized into close or open events using a segmental K-means (SKM) algorithm with a cut-off resolution of 70 μs. Clusters of single-channel bursts were selected by eye before being separated into lists of discrete bursts for every patch. *τ*_crit_ values were determined for each patch in order to preserve the 3 shortest close and 3 shortest open states. A linear, single-gate way 3C3O model scheme that displayed the largest ΣLL value for among both control and Shisa7 conditions was used for each patch. This particular arrangement has also been previously employed to model GABA_A_R activation (Dixon et al., 2014; Keramidas and Harrison, 2010; Lema and Auerbach, 2006). MIL estimation was used to obtain single-channel burst kinetic properties and dwell time distributions.

### Modeling and Simulation of Macroscopic Currents

The QuB software was used to simulate macroscopic GABA currents. Specifically, close and open states, along with corresponding rate constants between them were arranged in the aforementioned 3C3O model. Rate constants were set to fixed and a concentration-dependent binding step was appended to the second-to-last close state preceding channel opening (Figure 6A; purple arrow). To simulate the macroscopic GABA current, a protocol consisting of a single 1 ms application and 10 mM GABA was implemented. The number of channels simulated was set to 1000 and the response setting was set to deterministic. The simulation data files were exported to Clampfit 11.2 and mono- and bi-exponential fits were used to calculate the mean rise and weighted tau decay times, respectively.

### Statistical Analysis

Statistical analysis was performed using GraphPad Prism 9.0. All experiments were repeated at least three times. For concentration-response, log transformations of GABA concentrations were used and each data point was normalized to 10 mM GABA. Nonlinear, least squares regression using the log(agonist) and normalized response with variable slope was performed and the LogEC_50_ values between groups were compared. Shapiro-Wilk tests were used to assess normal distributions before between-group statistical comparisons. A two-tailed, Mann-Whitney U test was performed to assess for significant differences between control and Shisa7 conditions.

## Acknowledgements

We thank Dr. Joseph Lynch for providing us with the human β3 GABA_A_R subunit. We thank Drs. Gabriela Popescu and Gary Iacobucci for providing the QuB software. We thank Drs. Stefano Vicini, Marek Brodzki, and Gary Iacobucci for helpful discussions regarding kinetic analysis of single-channel data and stimulation of macroscopic currents. This work was supported by the NIH/NINDS Intramural Research Program (to W.L.)

## Author Contributions

D.C. and W.L. designed the project, and W.L. supervised the project. D.C performed electrophysiological recordings. D.C. and K.W performed cell-culture and preparation of single-channel data for kinetic analysis. D.C and A.K performed modeling and simulations of macroscopic currents. D.C. and W.L wrote the manuscript, and all authors read and commented on the manuscript.

## Declaration of Interests

The authors declare no competing interests.

## Notes

### Competing Interest Statement

The authors have declared no competing interest.

